# Linking alpha oscillations, attention and inhibitory control in adult ADHD with EEG neurofeedback

**DOI:** 10.1101/689398

**Authors:** Marie-Pierre Deiber, Roland Hasler, Julien Colin, Alexandre Dayer, Jean-Michel Aubry, Stéphanie Baggio, Nader Perroud, Tomas Ros

**Affiliations:** Division of Psychiatric Specialties, Department of Psychiatry, University Hospitals of Geneva, Geneva, Switzerland; Department of Psychiatry, University of Geneva, Geneva, Switzerland; Department of Basic Neurosciences, Geneva Medical Center, University of Geneva, Geneva, Switzerland; Division of Prison Health, University Hospitals of Geneva, Geneva, Switzerland; Department of Psychiatry, Dalhousie University, Halifax, NS, Canada

**Keywords:** adult ADHD, EEG, alpha oscillations, neurofeedback, inhibition control

## Abstract

Abnormal patterns of electrical oscillatory activity have been repeatedly described in adult ADHD. In particular, the alpha rhythm (8-12 Hz), known to be modulated during attention, has previously been considered as candidate biomarker for ADHD. In the present study, we asked adult ADHD patients to self-regulate their own alpha rhythm using neurofeedback (NFB), in order to examine the modulation of alpha oscillations on attentional performance and brain plasticity. Twenty-five adult ADHD patients and 22 healthy controls underwent a 64-channel EEG-recording at resting-state and during a Go/NoGo task, before and after a 30 min-NFB session designed to reduce (desynchronize) the power of the alpha rhythm. Alpha power was compared across conditions and groups, and the effects of NFB were statistically assessed by comparing behavioral and EEG measures pre-to-post NFB. Firstly, we found that relative alpha power was attenuated in our ADHD cohort compared to control subjects at baseline and across experimental conditions, suggesting a signature of cortical hyper-activation. Both groups demonstrated a significant and targeted reduction of alpha power during NFB. Interestingly, we observed a post-NFB increase in resting-state alpha (i.e. rebound) in the ADHD group, which restored alpha power towards levels of the normal population. Importantly, the degree of post-NFB alpha normalisation during the Go/NoGo task correlated with individual improvements in motor inhibition (i.e. reduced commission errors and slower reaction times in NoGo trials) only in the ADHD group. Overall, our findings offer novel supporting evidence implicating alpha oscillations in inhibitory control, as well as their potential role in the homeostatic regulation of cortical excitatory/inhibitory balance.

**Highlights:**

- Resting alpha power is reduced in adult ADHD suggesting cortical hyper-activation
- Adult ADHD patients successfully reduce alpha power during neurofeedback
- A post-neurofeedback rebound normalizes alpha power in adult ADHD
- Alpha power rebound correlates with improvement of inhibitory control in adult ADHD

## 1. Introduction

Attention-deficit hyperactivity disorder (ADHD) is characterized by symptoms of inattention and/or impulsivity and hyperactivity (Biederman and Faraone, 2005). While 2-7% of children are concerned worldwide (Sayal et al., 2018), the disorder often persists at later age with a prevalence of 4 to 5% in adulthood (Biederman and Faraone, 2005, Kessler et al., 2006, Sibley et al., 2017, Spencer et al., 2007). ADHD is associated with negative long-term outcomes such as impaired social adjustment, academic problems and high probability of psychiatric comorbid disorders (Gillberg et al., 2004, Katzman et al., 2017, Kessler et al., 2006, Klein et al., 2012, Skirrow and Asherson, 2013).

Historically, studies in children with ADHD initially observed generalized slowing of the EEG, characterized by an increase in slower frequency power (i.e., theta 4-7 Hz) and a reduction in faster frequency power (i.e, beta 14-25 Hz) (Arns et al., 2013). Given that the Theta/Beta Ratio (TBR) is known to decline during healthy development (Perone et al., 2018), an elevated TBR seen in ADHD has been proposed to reflect developmental delay and/or cortical hypoarousal (Barry et al., 2003, Sangal and Sangal, 2015). However, more recent studies have challenged the association of TBR with arousal, and its validity as a reliable discriminator of ADHD diagnosis (Arns et al., 2013, Lenartowicz and Loo, 2014). Furthermore, in some cases, elevated theta power can also reflect slowing of the alpha frequency, rather than reflecting an independent increase in theta power (Lansbergen et al., 2011). Hence, besides TBR, resting-state alpha power (8-12 Hz) has been the subject of several studies in adult ADHD patients. In healthy subjects, alpha power increases or decreases have been found to reflect cortical inhibition or excitation, respectively (Haegens et al., 2011, Klimesch, 2012, Mathewson et al., 2011, Romei et al., 2008). Functionally, elevated alpha amplitude has been associated with a diminished perception of sensory stimuli, and an internally-oriented state favoring mind wandering and attentional lapses (Mathewson et al., 2009, Ros et al., 2013, Sigala et al., 2014, van Dijk et al., 2008). In addition, increased alpha amplitude at recording sites overlying the motor cortex has been linked to voluntary motor inhibition (Hummel et al., 2002, Sauseng et al., 2013). While some ADHD studies have described an elevated level of alpha power compared to healthy controls (Bresnahan and Barry, 2002, Koehler et al., 2009, Poil et al., 2014), others have found a reduced level (Loo et al., 2009, Ponomarev et al., 2014, Woltering et al., 2012), or no significant differences (Bresnahan et al., 2006, Hermens et al., 2004, van Dongen-Boomsma et al., 2010). Hence, the contradictory alpha power results across studies may be viewed as supporting evidence for the possibility of multiple ADHD electrophysiological biotypes (Loo et al., 2018).

In the face of such heterogeneity, the use of brain-computer interfaces to provide real-time feedback (i.e. neurofeedback) in order to enable control of specific brain oscillations represents an intriguing option (see (Gruzelier, 2014) for a review). In this context, neurofeedback may be used to provide ADHD patients with instantaneous feedback of their EEG dynamics, in order to help them influence their own cortical activity and potentially improve various clinical features such as attention and inhibition (Niv, 2013, Sitaram et al., 2017). Here, select EEG parameters are converted into visual or auditory information and fed back in real time, allowing prolonged training with the goal of impacting brain plasticity (Ros et al., 2014, Sitaram et al., 2017). Neurofeedback-induced plasticity has been demonstrated in corticomotor (Ros et al., 2010) and cortico-striatal circuits (Koralek et al., 2012), which are relevant to the pathology of ADHD (Spencer et al., 2007). Importantly, emerging research has shown that neurofeedback may be used to improve inattention and impulsivity symptoms in ADHD (Arns et al., 2009, Micoulaud-Franchi et al., 2014), with some studies indicating effect-sizes close to that of methylphenidate and long-term impact of at least 6 months in adults (González-Castro et al., 2016, Mayer et al., 2016).

In light of consistent findings linking alpha-band changes with both attention and cortical inhibition (Lenartowicz et al., 2018), yet conflicting alpha-band signatures in ADHD, the present study aimed to reduce alpha power in adult ADHD patients during a single NFB session, in order to explore short-term plastic effects on omission (i.e. perceptual) and commission (i.e. motor inhibition) errors during a Go/NoGo task. Our principal hypotheses were based on existing evidence indicating that adults with ADHD exhibit increased baseline alpha power (Bresnahan and Barry, 2002, Koehler et al., 2009, Poil et al., 2014), which has separately been found to positively predict errors in response inhibition (Mazaheri et al., 2009). Specifically, our initial predictions were the following: 1) ADHD patients would exhibit significantly higher baseline alpha power compared to healthy controls; 2) ADHD patients would exhibit significant alpha power reductions post-NFB training, and 3) the degree of alpha reduction would predict improvements during Go/NoGo performance. As a control for patient-specific effects, a group of age-matched healthy participants were administered the same procedure. Here, we sought to answer whether different alpha power levels explained behavioral differences between ADHD patients and healthy controls, as well as, whether NFB had converging or diverging effects on alpha power and related behavioral performance in the two groups.

## 2. Material and Methods

### 2.1 Participants and experimental design

Twenty-five adult patients with ADHD (13 female, mean age: 33.9, SD: 10.9) were recruited in a specialized center for the assessment, treatment and care of patients suffering from ADHD at the Department of Psychiatry of the University Hospitals of Geneva. At the time of recruitment (usually several months after the initial contact with our center), 10 patients were unmedicated, 10 were taking methyphenidate, 2 atomoxetine, 1 antiepileptic, 1 benzodiazepine, 1 neuroleptic. Patients with comorbid psychiatric conditions were excluded. Twenty-two healthy adults (HC, 14 female, mean age: 31.1, SD: 7.4) were additionally recruited through announcements in the general population. Mean age between groups did not differ significantly (unpaired t-test, t = .996, p = .325). Prior to the study, written informed consent was obtained from each participant. The study was approved by the Research Ethic Committee of the Republic and Canton of Geneva [project number 2017-01029].

During a first clinical visit, patients and controls underwent three clinical questionnaires: (i) the ADHD Child Evaluation for Adults (ACE+), a semi-structured interview developed to support healthcare practitioners in the assessment and diagnosis of adults with ADHD (freely available at: https://www.psychology-services.uk.com/adhd.htm), (ii) the French version of the Structured Clinical Interview for DSM-IV Axis II Personality Disorders (SCID-II, (First et al., 1997)) and (iii) the French version of the Diagnostic Interview for Genetic Studies (DIGS, mood disorder parts only (Preisig et al., 1999)). Exclusion criteria included: history of head injury with loss of consciousness, epilepsy or stroke, non-neurological conditions susceptible to impair brain function (e.g., cancer or cardiovascular disease), and other current psychiatric disorders based on the above mentioned semi-structured interviews: major depressive disorder, bipolar disorder, anxiety disorders, borderline personality disorder, and substance use disorders. All patients treated with psychostimulants stopped their medication 24 h before the experimental visit. Among the 25 patients, 18 were of mixed subtype, 6 of inattentive subtype, and 1 of hyperactive subtype.

### 2.2 EEG procedure

The experiment, designed to evaluate the effect of 30 min NFB session on EEG at rest with eyes opened (EO) and during performance of a Continuous Performance Task (CPT), consisted in three sequential parts: EEG-evaluation 1, EEG-NFB, and EEG-evaluation 2 (Fig. 1). A 3 min baseline resting state with eyes opened (EO1) preceded EEG-evaluation 1, which consisted of (i) 6 min of the CPT (CPT1), (ii) self-rated questionnaires assessing instantaneous state anxiety and arousal, and (iii) 3 min of EO rest (EO2). Then, the subject underwent 30 min of EEG-NFB session, as detailed below. Lastly, EEG-evaluation 2 consisted of (i) 3 min of EO rest (EO3), (ii) self-rated questionnaires assessing instantaneous state anxiety and arousal, and (iii) 6 min of the CPT (CPT2). The CPT consisted in the sequential presentation of 16 letters for 200 ms. The subjects were asked to press the left mouse button when any letter except the target letter “X” appeared. There was a total of 240 trials, with 75% Go trials and 25% NoGo trials. The maximal response window was of 600 ms, with a varying intertrial interval (800, 900 or 1000 ms). The self-rated questionnaires were the state anxiety part of the Spielberger’s State Anxiety Inventory (STAI) and the Thayer’s Activation-Deactivation Adjective Checklist. Questionnaire data were not reported here to avoid result section overload.

**Figure 1.**
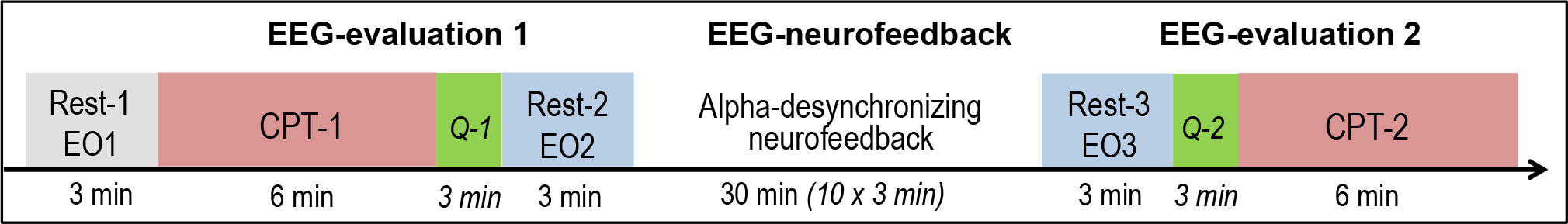
Timeline of the experimental procedure. EO: eyes opened; CPT: Continuous Performance Task; Q: self-rated questionnaires (Spielberger’s and Thayer’s).

EEG was recorded continuously using 64 Ag/AgCl electrode cap according to the 10-20 international system, with a sampling rate of 500 Hz. The ground electrode was placed on the scalp at a site equidistant between Fpz and Fz, and the reference electrode at CPz. Electrical signals were amplified using the Eego mylab system (ANT Neuro, Netherlands), and all electrode impedances were kept below 5 k?. For offline analyses, EEG signals were re-referenced to common-average reference. Independent component analysis (ICA) was used to identify and remove stereotypical artifacts using the Infomax algorithm (blinking and lateral eye movements) (Jung et al., 2000). Statistically defined artifact rejection was then carried out with the FASTER method (Nolan et al., 2010) removing segments based on extreme deviations of amplitude and variance from the mean.

### 2.3 Neurofeedback procedure

The EEG neurofeedback training protocol is fully described elsewhere (Kluetsch et al., 2014, Ros et al., 2013). Briefly, the Pz channel was specifically used for neurofeedback, using a Pro-Comp amplifier interfacing with EEGer 4.2 neurofeedback software (EEG Spectrum Systems, CA). Separate ground and reference electrodes were placed at on the right and left earlobe, respectively. Pz was selected as the electrode overlying the posterior parietal cortex, whose metabolic changes have been previously linked to EEG alpha rhythm modulation (Laufs et al., 2006). All participants interacted with a ‘SpaceRace’ game where they received continuous visual feedback in the form of a moving spaceship and a dynamic bar graph whose height was inversely proportional to real-time alpha amplitude fluctuations. Participants were told that the spaceship would move forward whenever they were ‘in-the-zone’ of their target brain activity (i.e., alpha lower than threshold), and that it would stop when they were ‘out-of-the-zone’ (i.e., alpha higher than threshold). The aim of the training was to use the feedback they received during the game to learn to keep the spaceship traveling through space. For the purpose of online neurofeedback training, the EEG signal was infinite impulse response band-pass filtered to extract alpha (8-12 Hz) with an epoch size of 0.5 s. Participants were rewarded upon suppression of their absolute alpha amplitude. For each participant, the reward threshold was initially set so that their alpha amplitude would fluctuate below the initial 3-min baseline average approximately 60% of the time (i.e., they received negative feedback about 40% of the time). To ensure that all participants received comparable frequencies of reward, we readjusted their reward thresholds to meet the desired ratio, when they achieved disproportionately higher (> 80%) or lower (< 40%) rates of reward during feedback. The entire neurofeedback session was divided into 3-min training periods with a short break (10 s) after each period. During the breaks, the scores for the preceding periods were displayed.

### 2.4 Data analysis

#### 2.4.1 Alpha spectral power in the 6 conditions

EEG spectrum was obtained using Brain Vision Analyzer 2 (Brain Products GmbH) via Fast Fourier Transform (FFT) on 2048 ms non-overlapping Hanning-windowed epochs, allowing a frequency resolution of 0.5 Hz. Relative alpha power was calculated in the 8-12 Hz bandwidth (reflective of the NFB protocol) as the absolute alpha power divided by to the full spectrum power (1.5 to 40 Hz). The mean relative alpha power was computed across the 64 electrodes. A repeated-measures ANOVA with 6-level Condition (EO1, CPT1, EO2, NFB, EO3, CPT2) as within-subject, and 2-level Group (ADHD, HC) as between-subject factors was used to evaluate statistical differences of the mean relative alpha power between groups and conditions. For the ANOVA, Huynh–Feldt correction for non-sphericity was applied when appropriate. Topographical analysis of EEG spectral data were further carried out with the Neurophysiological Biomarker Toolbox (NBT, http://www.nbtwiki.net/) in Matlab (MathWorks Inc.), after 0.5 to 40 Hz band-pass filtering and a 55-65 Hz notch filter. To test for group/condition differences, we used a permutation test with 5000 repetitions (Nichols and Holmes, 2002) on all channels, and subsequently corrected for multiple comparisons using binomial correction (Poil et al., 2014). The significance threshold for all comparisons was set to alpha = 0.05.

#### 2.4.2 Alpha event-related desynchronization (ERD) during CPTs

For analysis of event-related EEG oscillations, the EEG was segmented per trial type (Go and NoGo) in both CPT conditions into epochs of 1900 ms, starting 800 ms before stimulus onset. Only trials corresponding to correct responses were considered. A time-frequency analysis based on a continuum wavelet transform of the signal (complex Morlet’s wavelets) was applied to each epoch from 1 to 30 Hz in 1-Hz steps (Tallon-Baudry et al., 1998). The resulting dataset consisted in an average TF representation of the signal over all trials of the same type. Using Matlab scripts, we extracted the time course of the event-related desynchronization/synchronization (ERD/ERS) in the 8-12 Hz frequency range for each participant, relative to a baseline calculated between 800 and 100 ms before stimulus onset. Mean alpha ERD amplitude was calculated for each electrode. According to the topographic distribution of the alpha ERD, we computed the mean alpha ERD over the 28 posterior electrodes, including centro-parietal, parietal, parieto-occipital and occipital channels. For each trial type (Go, NoGo), a repeated-measures ANOVA with 2-level Condition (CPT1, CPT2) as within-subject, and 2-level Group (ADHD, HC) as between-subject factors was used to evaluate statistical differences of the mean alpha ERD amplitude between groups and conditions. Statistical threshold was set at p < .05 after Huynh–Feldt correction for non-sphericity when appropriate. Post-hoc analysis used paired t-tests with p < .05 as significance threshold after Bonferroni correction for multiple comparisons.

#### 2.4.3 Performance at CPTs

Errors included omissions (missed targets) and commissions (responses to non-targets or false alarms). D-prime was defined by the ratio between hits (correct responses) and commissions (false alarms), providing a measure of stimulus discriminability. Reaction time (RT) corresponded to the time interval between stimulus onset and mouse button press. RT variability (SD RT) and RT variation coefficient (Var RT), which provide information on the variability of RT, were also examined. Perseveration, defined as response with a RT < 150 ms, was discarded because it lacked of variance. A repeated-measures ANOVA with 2-level Condition (CPT1, CPT2) as within-subject, and 2-level Group (ADHD, HC) as between-subject factors was used to evaluate statistical differences of performance between groups and conditions. Statistical threshold was set at p < .05 after Huynh–Feldt correction for non-sphericity when appropriate. Post-hoc analysis used paired t-tests with p < .05 as significance threshold after Bonferroni correction for multiple comparisons.

#### 2.4.4 Correlation analyses

To examine the relation between electrophysiological activities and behavioral/mood parameters, as well as their modulation by NFB training, we calculated the absolute differences between CPT2 and CPT1 of the respective measures and computed the following correlations in each group (Pearson coefficient): i) CPT2-CPT1 relative alpha power vs CPT2-CPT1 performance parameters; ii) CPT2-CPT1 alpha ERD vs CPT2-CPT1 performance parameters; iii) CPT2-CPT1 relative alpha power vs CPT2-CPT1 alpha ERD.

Statistical analyses were conducted with SPSS 25.

## 3. Results

### 3.1 Alpha power between groups, and across the 6 conditions

Fig. 2 presents the mean relative alpha power value in the 6 conditions for the ADHD and HC groups. There was a significant Group effect on alpha power across the 6 conditions (F= 4.10, p < .05), ADHD displaying lower alpha power than HC. A significant Condition effect (F = 32.20, p < .001) and a significant Group x Condition interaction (F = 3.41, p < .05) were observed.

**Figure 2.**
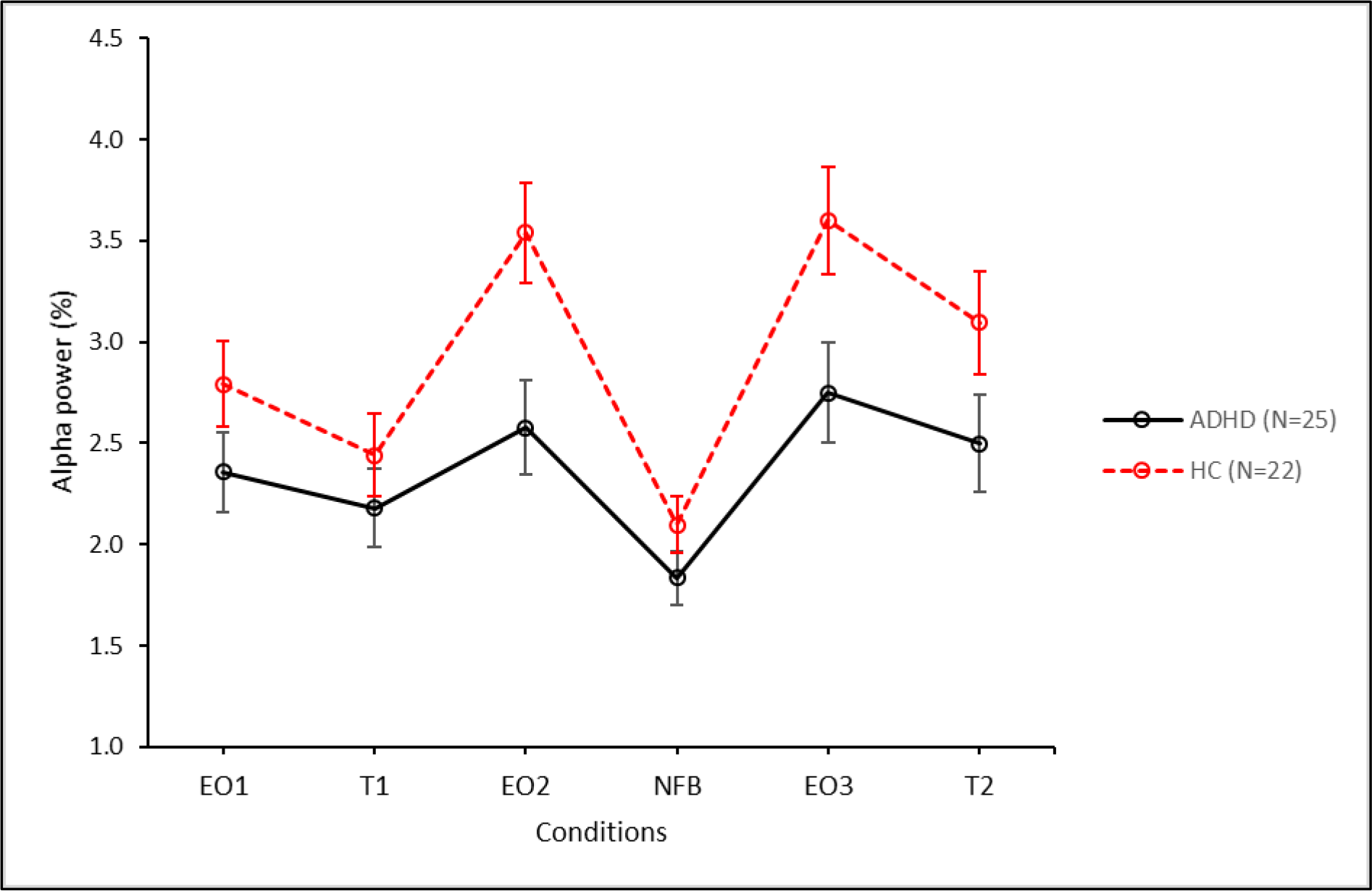
Relative alpha power for ADHD (black solid line) and healthy controls HC (red dashed line) in each condition (average over the 64 electrodes). Bars represent confidence intervals.

In a nutshell, selected contrasts were performed in accordance with our *a priori* hypotheses:

(A) comparison of relative alpha power at baseline (EO1) between ADHD and HC; (B) evaluation of NFB effect on relative alpha power in ADHD and HC respectively (NFB vs EO2); (C) comparison of relative alpha power at rest pre-and post-NFB in ADHD and HC respectively (EO3 vs EO2); (D) comparison of relative alpha power during CPT pre-and post-NFB in ADHD and HC respectively (CPT2 vs CPT1).

#### A. Resting-state EEG differences between ADHD patients and control subjects

The relative alpha power was significantly lower in ADHD patients than in healthy controls (HC) at baseline resting state (EO1) in the frontal region (binomial corrected, p < .05) (Fig. 3).

**Figure 3.**
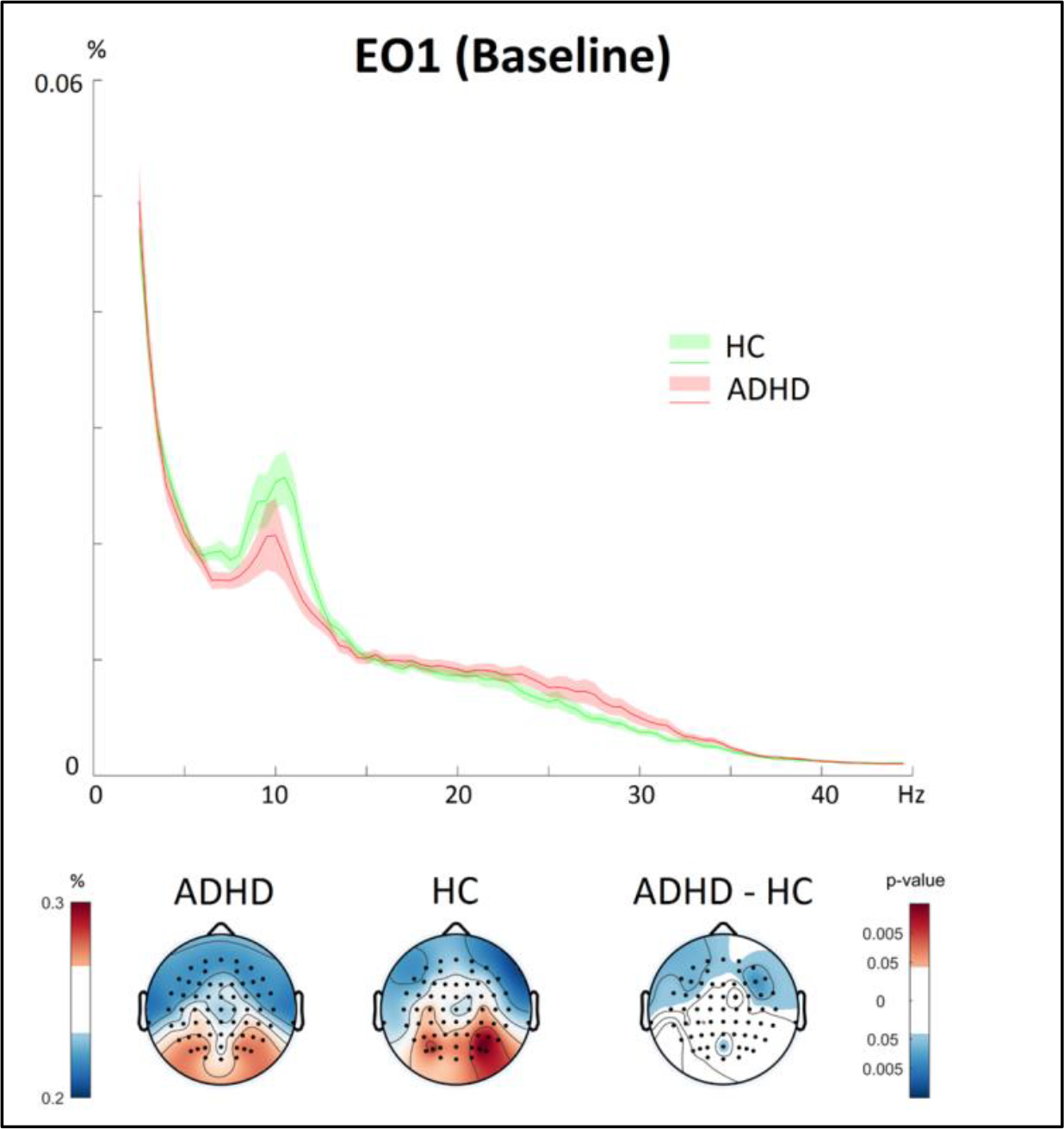
Top: EEG relative power spectrum at baseline (EO1) in ADHD patients (red) and healthy subjects (HC, green). Solid lines: mean relative value over the 64 electrodes, highlighted areas: confidence interval. Bottom: Topographic plots of relative alpha amplitude in EO1 for the ADHD and HC groups, and unpaired permutation test (binomial corrected, p < .05).

#### B. EEG signatures during neurofeedback training

Relative alpha power was successfully reduced during NFB as compared to EO2 in both groups (NFB – EO2, binomial corrected, p < .05), attesting that independently of diagnosis, the participants successfully downregulated their alpha amplitude (Fig. 4).

**Figure 4.**
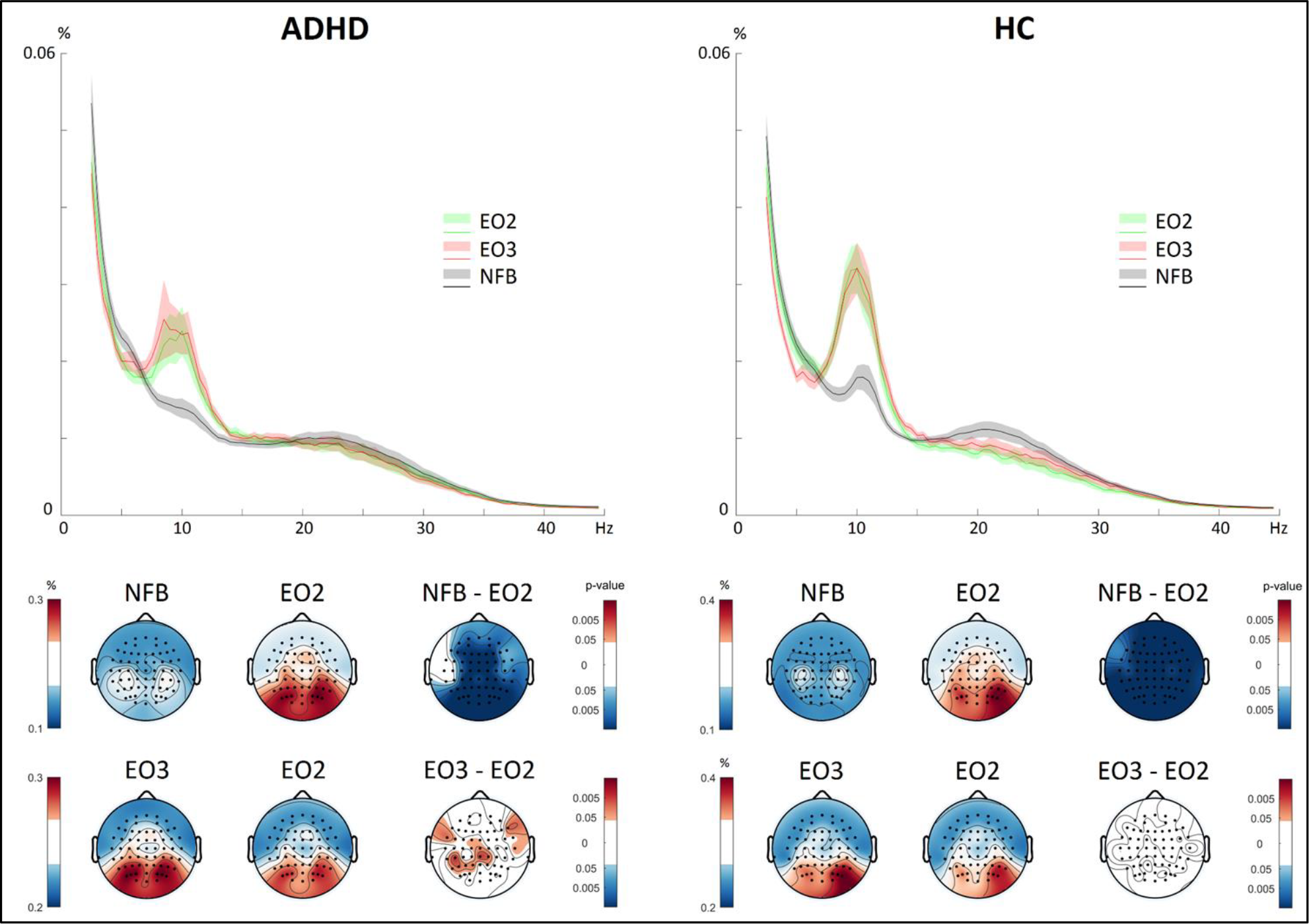
Top: EEG relative power spectrum during EO2 (green), EO3 (red) and NFB (grey) in ADHD (left) and HC (right). Solid lines: mean relative value over the 64 electrodes, highlighted areas: confidence interval. Bottom, first row: Topographic plots of relative alpha amplitude in NFB and EO2, and paired permutation test (binomial corrected, p < .05). Bottom, second row: Topographic plots of relative alpha amplitude in EO3 and EO2, and paired permutation test (binomial corrected, p < .05).

#### C. Resting-state EEG signatures pre-to-post neurofeedback

As shown in Fig. 4, a significant rebound of alpha power post-NFB (EO3) as compared to pre-NFB (EO2) was evident only for the ADHD group (binomial corrected, p < .05).

#### D. Continuous Performance Test EEG signatures pre-to-post neurofeedback

As depicted in Fig. 5, comparing the CPT EEG pre-to post-NFB revealed higher alpha power post-NFB (CPT2) as compared to pre-NFB (CPT1) in both groups (binomial corrected, p < .05). This indicates different levels of alpha in the same individuals during the CPT task, pre-to-post NFB.

**Figure 5.**
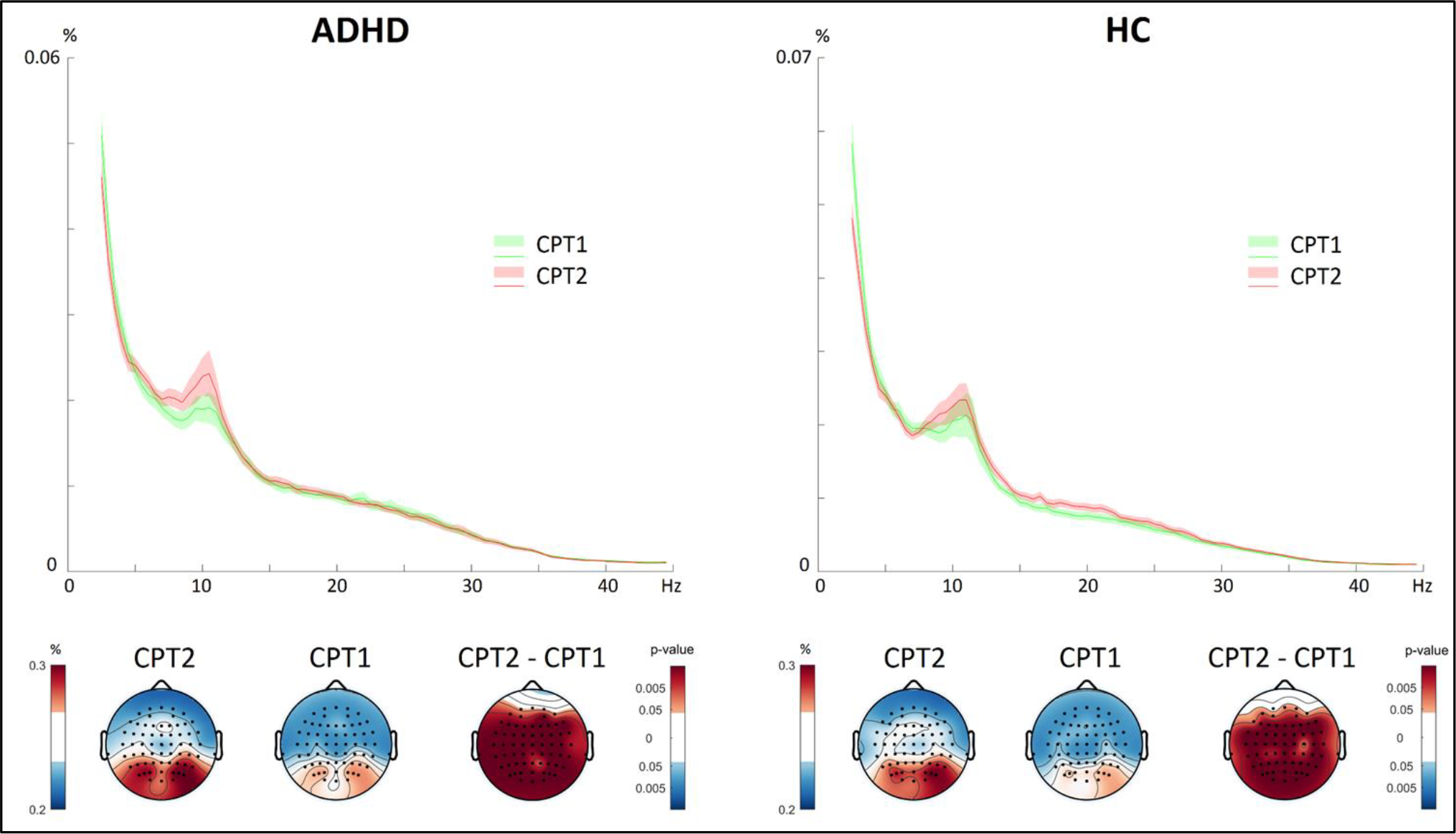
Top: EEG relative power spectrum during CPT1 (green) and CPT2 (red) in ADHD (left) and HC (right). Solid lines: mean relative value over the 64 electrodes, highlighted areas: confidence interval. Bottom: Topographic plots of relative alpha amplitude in CPT2 and CPT1, and paired permutation test (binomial corrected, p < .05).

### 3.2 Alpha event-related desynchronization (ERD) in CPT Go and NoGo trials, pre-and post-NFB

For both Go and NoGo trials, there was a significant condition effect but no significant group effect, nor significant group x condition interaction, on the mean alpha ERD amplitude (Go trials, F = 23.00, p < .001; NoGo trials, F = 20.54, p < .001). Overall in both groups and both trial types, the alpha ERD was larger in CPT2 than CPT1 (see Supplementary Table 1 in Appendix).

### 3.3 CPT performance pre-and post-NFB

There was a significant group effect on the following CPT parameters: omission (F = 10.40, p < .01), commission (F = 12.83, p < .001), d-prime (F = 25.50, p < .001), SD RT (F = 16.23, p < .001), Var RT (F = 27.09, p < .001). This indicates that, compared to the HC and independently of the pre-or post-NFB condition, the ADHD group committed more omission (i.e. detection) and commission (i.e. motor inhibition) errors, and demonstrated more variability in RT. Additionally, there was a significant condition effect on d-prime (F = 5.46, p < .05), stimulus detectability being higher post-NFB, and on Var RT (F = 6.46, p < .05), RT variability being reduced post-NFB (see Supplementary Table 1 in Appendix).

### 3.4 Correlation analyses

#### 3.4.1 Alpha power and CPT performance pre-and post-NFB (CPT2-CPT1)

In the ADHD group, there was a significant negative correlation between CPT2-CPT1 relative alpha power and CPT2-CPT1 commission errors (r = -.483, p < .05), so that the larger the alpha rebound (i.e. increase) at CPT2, the less commission errors were committed (Fig. 6A). There was also a significant positive correlation between CPT2-CPT1 relative alpha power and CPT2-CPT1 reaction time (r = .471, p < .05). Hence, the larger the alpha rebound at CPT2, the slower was the reaction time.

No significant correlations were found in the HC group.

**Figure 6.**
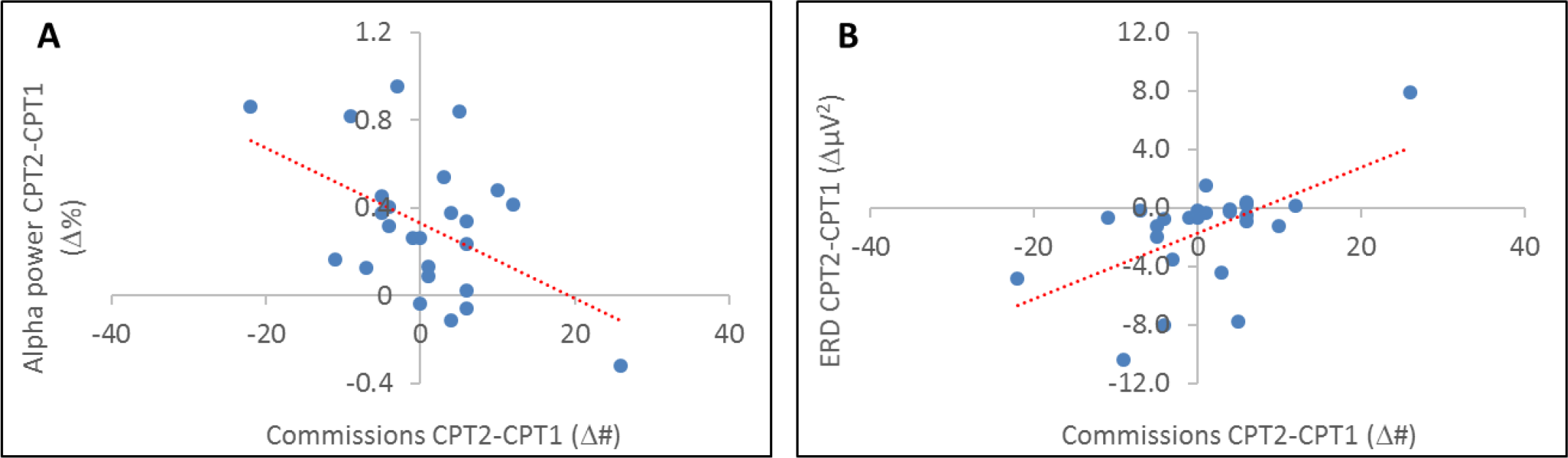
Correlation between commission differences and A. CPT2-CPT1 relative alpha power differences, B. CPT2-CPT1 NoGo trials ERD differences in ADHD.

#### 3.4.2 Alpha ERD and CPT performance pre-and post-NFB (CPT2-CPT1)

In the ADHD group, there was a significant positive correlation between CPT2-CPT1 alpha ERD amplitude and CPT2-CPT1 commission errors (Go trials: r = .527, p < .01; NoGo trials: r = .568, p < .01), so that the greater the alpha ERD amplitude at CPT2 (negatively) in Go and NoGo trials, the less commission errors were committed (Fig. 6B). There was also a significant negative correlation between CPT2-CPT1 Go alpha ERD amplitude and CPT2-CPT1 reaction time (r = -.404, p < .05), so that the largest was the Go alpha ERD amplitude at CPT2 (negatively), the slower was the reaction time.

No significant correlations were found in the HC group.

#### 3.4.3 Alpha power and alpha ERD

In both groups, there was a significant negative correlation between CPT2-CPT1 relative alpha power and CPT2-CPT1 alpha ERD amplitude in Go and NoGo trials (ADHD: Go trials, r= -.828, p < .001, NoGo trials, r= -.782, p < .001; HC: Go trials, r= -.685, p < .001, NoGo trials, r= -.653, p < .001). Hence, the largest was the alpha rebound at CPT2, the largest was the alpha ERD amplitude at CPT2 (negatively).

#### 3.5 Theoretical model of neurofeedback effects: homeostatic normalisation of excitation/inhibition (E/I) in ADHD (Fig. 7)

Experiments in humans and animals have firmly established that brain activity and E/I balance are homeostatically regulated, where intrinsic mechanisms exist to limit neural excitability or neuronal firing from reaching abnormally high/low extremes, in order to preserve neural network function (Karabanov et al., 2015, Maffei and Fontanini, 2009). Prevailing models of the alpha rhythm have proposed that it acts as an “inhibitory gate” for sensorimotor cortices (Jensen and Mazaheri, 2010), and therefore alpha power may be considered to inversely correlate with E/I balance. Accordingly, alpha oscillations display a negative correlation with cortical activation (Podvalny et al., 2015) and metabolism (Conner et al., 2011). Thus, a signature of abnormally reduced alpha power as shown by our cohort with ADHD would indicate a state of increased E/I, while the ‘high-alpha’ biotype (Bresnahan and Barry, 2002, Koehler et al., 2009, Poil et al., 2014) would reflect low E/I. The proposed U model in Fig. 7 indicates that normalizing alpha power (and therefore E/I balance) towards healthy population values would improve inhibitory performance for both ‘high’ and ‘low’ alpha biotypes.

**Fig 7.**
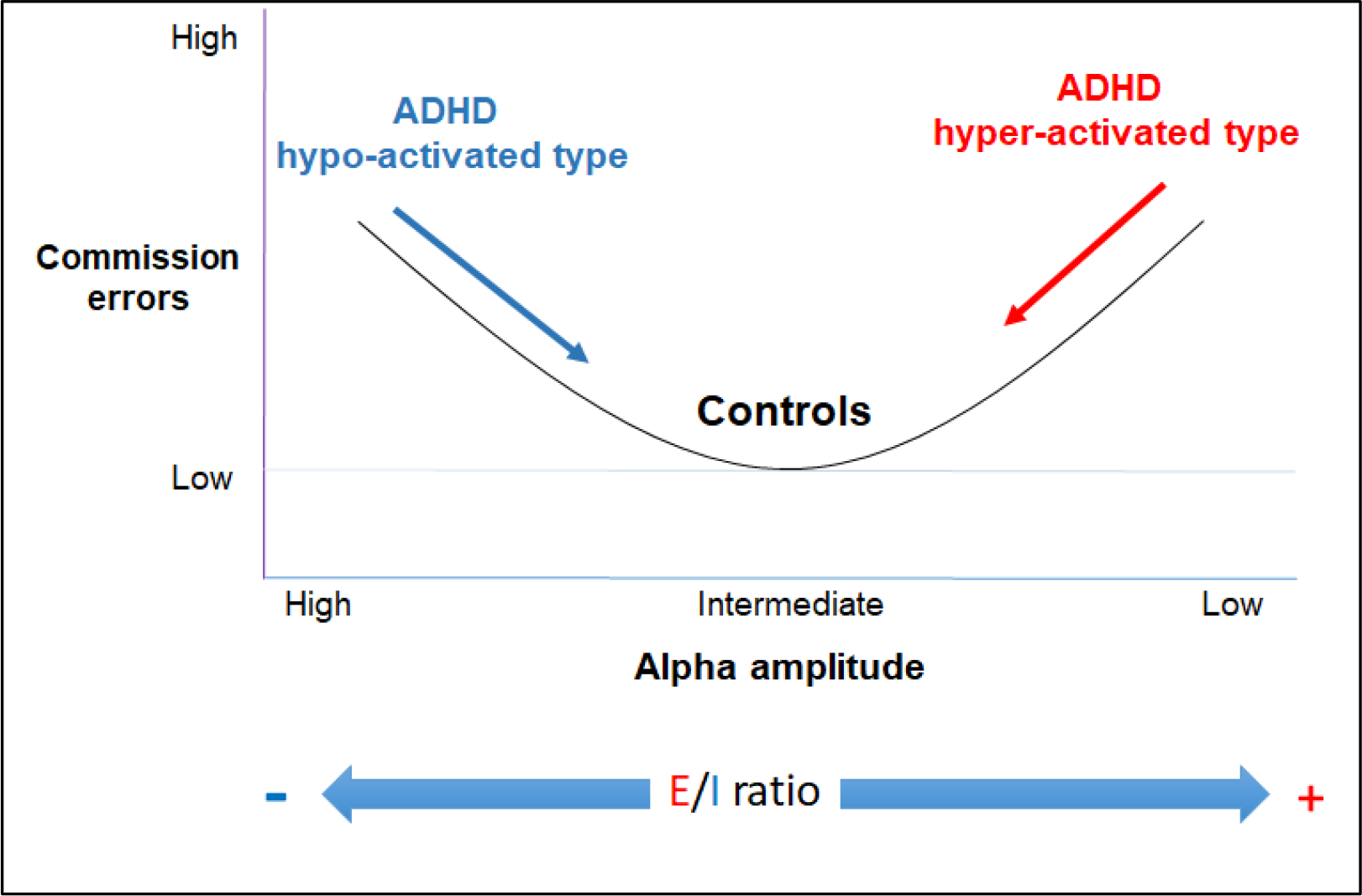
Theoretical model linking alpha oscillations, cortical E/I balance and behavioral performance. U-relationship between elevated response inhibition (i.e. commission) errors, alpha amplitude and cortically hyper-or hypo-activated ADHD biotypes.

## 4. Discussion

The present study focused on the relationship between alpha oscillations, attention, and motor inhibition in adult ADHD, using an experimental design with resting and task conditions, including a single neurofeedback session designed to modulate within-subject alpha power. Firstly, at baseline resting state, adults with ADHD exhibited lower relative alpha power than healthy control subjects, suggesting higher levels of cortical activation (Conner et al., 2011, Podvalny et al., 2015), and in line with a ‘low-alpha’ biotype (Loo et al., 2009, Ponomarev et al., 2014, Woltering et al., 2012). Secondly, consistent with studies in other populations (Kluetsch et al., 2014, Ros et al., 2013), we demonstrated for the first time that adult ADHD patients successfully downregulated their alpha rhythm during neurofeedback, and to a similar degree as control subjects. Thirdly, a significant increase (termed ‘rebound’) of post-NFB resting alpha power was observed in ADHD patients, partially restoring alpha power towards baseline levels seen in control subjects. Interestingly, increased post-NFB alpha power during the CPT correlated with improvements in motor inhibition in ADHD patients only.

### 4.1 Resting-state alpha power in adult ADHD

Contrary to our initial predictions, resting state alpha power in our adult ADHD sample was significantly reduced compared to control subjects. Hence, the signature of our cohort of adult ADHD patients was more consistent with a ‘low-alpha’ biotype (Loo et al., 2009, Ponomarev et al., 2014, Woltering et al., 2012) rather than the ‘high-alpha’ biotype (Bresnahan and Barry, 2002, Koehler et al., 2009, Poil et al., 2014). Thus far, no consistent pattern has emerged from studies investigating alpha-band spectral power at rest in adult ADHD. In line with the ‘EEG slowing’ signature in childhood ADHD, represented by elevated power of low-frequency rhythms (e.g. delta, theta) (Clarke Adam R. et al., 2001), there have been several reports of elevated power in the dominant low-frequency (i.e. alpha) rhythm in adult ADHD (Bresnahan and Barry, 2002, Koehler et al., 2009, Poil et al., 2014). Alpha oscillations have been associated with reduced excitability and neuronal firing of sensory cortices (Haegens et al., 2011, Romei et al., 2008), functionally ‘gating’ access to external sensory stimuli (Jensen and Mazaheri, 2010, Macdonald et al., 2011). However, the view that ADHD is merely an underarousal disorder has begun to be challenged by various studies reporting excess high-frequency beta rhythms in some subtypes of ADHD (Clarke et al., 2011, Clarke A. R. et al., 2001, Loo et al., 2009, Meier et al., 2014). Importantly, this beta-rhythm biotype also displays reduced alpha power (Loo et al., 2009), an observation confirmed by the present study as well as other independent groups (Ponomarev et al., 2014, Woltering et al., 2012). The reduction of posterior alpha power has been associated with states of increased activation of visual areas (Ergenoglu et al., 2004, Hanslmayr et al., 2007, Romei et al., 2008, Sigala et al., 2014), coinciding with higher cortical metabolism (Conner et al., 2011, Laufs et al., 2006).

### 4.2 EEG signatures related to NFB training

The present study demonstrated successful NFB-related alpha desynchronization (i.e. reduction) in both ADHD and control groups. Interestingly, in spite of their lower baseline resting alpha power, ADHD patients succeeded in further desynchronizing their alpha rhythm through NFB, in accordance with previous work demonstrating bidirectional control of alpha oscillations (Ros et al., 2013, Zoefel et al., 2011). Alpha desynchronization is a cortically ‘activating’ form of NFB which has shown to have neurobehavioral effects in healthy subjects as well as patients with post-traumatic stress disorder (PTSD) (Kluetsch et al., 2014, Ros et al., 2010, Ros et al., 2013).

Following NFB, there was a significant alpha resynchronization in both groups. This could be interpreted as a form of homeostatic regulation, compatible with intrinsic mechanisms limiting neural excitability or neuronal firing from reaching abnormally high/low extremes (Davis, 2013, Karabanov et al., 2015, Turrigiano and Nelson, 2004). As a result, we show for the first time in adult ADHD a significant increase (termed ‘rebound’) of resting alpha power after NFB, in the direction of healthy control values. Interestingly, this almost exactly mirrors the effect of this NFB protocol in PTSD patients (Kluetsch et al., 2014) who also display lower-than-normal levels of resting alpha power (Ros et al., 2017), consistent with NFB models of homeostatic regulation (see (Ros et al., 2014) for a review)

### 4.3 EEG signatures during the Go/NoGo task

In terms of Go/NoGo errors, ADHD patients performed overall worse than control subjects at the CPT. As previously reported, they committed more omission (Go) and commission (NoGo) errors (Fasmer et al., 2016, Woltering et al., 2012) and were more variable in their reaction time (Fasmer et al., 2016, Kofler et al., 2013). Notably, independently of diagnosis, stimulus detectability significantly improved and reaction time variability reduced post-NFB, suggesting some improvement in perceptive and motor action processes. Critically, in ADHD patients only, both pre-to-post increases in alpha power and alpha ERD during the CPT predicted intra-individual reductions in commission errors as well as reaction time slowing, indicating task-related alpha power and alpha ERD as significant mediators of inhibitory control (Klimesch et al., 2007, Rihs et al., 2007). Hence, normalisation of E/I balance through alpha self-regulation could be directly responsible for improvements of motor inhibition in ADHD.

The electrophysiological abnormalities in ADHD suggest that (at least) along one dimension, the symptoms of ADHD could be explained by positive (over-activated) or negative (under-activated) deviations from an optimal E/I balance. Compatible with observations that ADHD appears to be electrophysiologically heterogeneous (Clarke et al., 2011), individual patients may be displaying patterns that reflect both cortical hypo-arousal and hyper-arousal, possibly residing on opposite sides of a U-curve (see Fig. 7 for a proposed model). Considering the results in Fig. 6 from this framework, and noting that our adult ADHD patients had abnormally reduced alpha power at baseline (i.e. high E/I), it is conceivable that this EEG profile might respond homeostatically to any extra increases in excitation (i.e. through NFB alpha reduction). If so, the correction by the system to reduce the ramping up of excessively high E/I might then manifests in an altogether opposite signature (i.e. increased alpha power). According to this framework, medium resting-state EEG power might coincide with optimal behavioral performance (Ros et al., 2014), as suggested by attentional deficits during extremes of high and low prefrontal activity (Pezze et al., 2014).

### 4.4 Task-related alpha ERD in adult ADHD

In response to visual stimuli, an alpha ERD is classically recorded over the parieto-occipital regions actively engaged in the attentional processing of visual information (Deiber et al., 2012, Klimesch, 2012, Rihs et al., 2007, Thut et al., 2006). We firstly observed that the alpha ERD was larger in both groups during the post-than pre-NFB CPT. Secondly, alpha ERD changes were negatively correlated with the alpha power difference between the two tasks, so that the larger the alpha power amplitude difference, the larger (i.e. the more negative) was the alpha ERD. This observation illustrates the reactivity of the alpha rhythm, potentially restoring its dynamic range, where stronger spontaneous alpha power provides a larger range for its desynchronization driven by an event (i.e the ERD) (Mayhew et al., 2013, Ros et al., 2014).

### 4.5 Limitations

We acknowledge limitations related to our study. Our sample size was small, limiting statistical power and the strength of our conclusions. Moreover, as both groups improved in signal detectability post-NFB, we did not observe a NFB-specific improvement in the ADHD compared to the control group. Our novel results require replication using larger patient and control samples, preferably including a ‘high-alpha’ ADHD biotype, in order to fully test the E/I normalisation model proposed in Fig 7.

### 4.6 Conclusions

Our results support studies of adult ADHD patients exhibiting cortical hyper-activation, as evidenced by low resting alpha power compared to healthy controls. Despite their reduced baseline alpha power, ADHD patients succeeded in further reducing their alpha rhythm during NFB, which was followed by a resynchronisation of resting-state alpha activity, compatible with models of homeostatic regulation of E/I balance. Notably, analysis of alpha power and ERD during performance of a Go/NoGo task revealed a significant association between post-NFB alpha power/ERD increase and improvements in motor inhibition (i.e. commission errors), supporting a key role of alpha oscillations in ADHD inhibitory deficits.

## Appendix

**Supplementary Table 1.** Performance parameters and alpha ERD amplitude pre (1) and post (2) NFB in both groups. Mean values with standard errors and p-values of pairwise comparisons between groups and between conditions (Bonferroni corrected). *: p < .05; **: p < .01; ***: p < .001; ns: non-significant.

| Parameter | Group | CPT | Mean ± se | Post-hoc<br>Group | comparisons<br>Condition |
| --- | --- | --- | --- | --- | --- |
| Omission (N) | ADHD | 1 | 6.68 ± 1.21 | p= .002** | p= .265 ns |
| | | 2 | 5.76 ± 1.38 | | |
| | CONTROL | 1 | 1.27 ± 1.29 | | |
| | | 2 | 0.27 ± 1.47 | | |
| Commission (N) | ADHD | 1 | 16.32 ± 1.61 | p< .001*** | p= .592 ns |
| | | 2 | 17.08 ± 1.38 | | |
| | CONTROL | 1 | 10.82 ± 1.71 | | |
| | | 2 | 8.91 ± 1.47 | | |
| d-prime | ADHD | 1 | 2.78 ± 0.17 | p< .001*** | p=.024* |
| | | 2 | 2.88 ± 0.17 | | |
| | CONTROL | 1 | 3.73 ± 0.18 | | |
| | | 2 | 4.16 ± 0.18 | | |
| Mean RT (ms) | ADHD | 1 | 328.49 ± 9.14 | p= .488 ns | p= .852 ns |
| | | 2 | 336.92 ± 8.98 | | |
| | CONTROL | 1 | 343.95 ± 9.74 | | |
| | | 2 | 337.84 ± 9.57 | | |
| SD RT (ms) | ADHD | 1 | 83.53 ± 3.21 | p< .001*** | p= .234 ns |
| | | 2 | 82.87 ± 3.43 | | |
| | CONTROL | 1 | 68.55 ± 3.42 | | |
| | | 2 | 63.58 ± 3.65 | | |
| Var RT (%) | ADHD | 1 | 25.45 ± 0.81 | p< .001*** | p= .015* |
| | | 2 | 24.32 ± 0.75 | | |
| | CONTROL | 1 | 20.08 ± 0.85 | | |
| | | 2 | 18.91 ± 0.78 | | |
| ERD Amplitude Go<br>(μV) | ADHD | 1 | -3.67 ± 1.11 | p= .504 ns | p< .001*** |
| | | 2 | -5.28 ± 1.44 | | |
| | CONTROL | 1 | -4.54 ± 1.18 | | |
| | | 2 | -6.88 ± 1.53 | | |
| ERD Amplitude<br>NoGo<br>(μV) | ADHD | 1 | -3.85 ± 1.15 | p= .486 ns | p< .001*** |
| | | 2 | -5.38 ± 1.47 | | |
| | CONTROL | 1 | -4.45 ± 1.23 | | |
| | | 2 | -7.39 ± 1.57 | | |

